# Neural multiplexing vs place coding: Multiplexing steps in when most needed

**DOI:** 10.64898/2026.08.06.743261

**Authors:** Julia M. Leeman, Shawn M. Willett, Nicholas Marco, Surya T. Tokdar, Jennifer M. Groh

## Abstract

Sensory scenes contain many different stimuli. Two complementary theories about how the brain segregates signals from different stimuli concern (a) time division multiplexing, such that neurons switch between encoding each item over time; and/or (b) place coding, such that different populations of neurons encode each item. Such time division multiplexing would appear to be required when the population of neurons responsive to each stimulus overlaps, as place coding lacks the granularity to resolve the two stimuli. This predicts that as responses to component stimuli become more similar, and thus less well resolved by place coding, there would be a greater incidence of multiplexing.

We tested this hypothesis using single-unit responses in the macaque inferior colliculus to combinations of two sounds of varying center frequencies (given that sound frequency is place coded in this structure). We found that neurons were more likely to multiplex when their responses to each individual sound was more similar, differentiating signals whose neural representations would otherwise be less distinct. This finding supports the theory that neurons multiplex to maintain information about concurrent stimuli when place coding is insufficient to prevent largely overlapping responses in the neural population.

## Introduction

Natural scenes contain many stimuli. The presence of multiple stimuli presents challenges to neural coding schemes. The problem is most apparent in the case of monotonic rate coding. If neurons encode a given stimulus parameter via a firing rate proportional to that parameter, it is not clear how the values for more than one stimulus could be encoded at the same time. For example, in the inferior colliculus (IC), horizontal sound location is encoded via firing rates that vary monotonically as a function of the azimuthal angle of the sound, typically reaching their maximal values for sounds lying along the axis of the contralateral ear (Groh et al., 2003; McAlpine & Grothe, 2003; Porter & Groh, 2006). If there are sounds at more than one location, it is uncertain how such a code can represent each of the sounds.

Two principal solutions to this problem have been proposed: a) that signals related to each stimulus can be “multiplexed” or interleaved across time; or b) that place coding along some other stimulus dimension could serve to sort the different signals into distinct neural populations, alleviating the need for multiplexing across time. In the case of the inferior colliculus, while sound location is rate coded, sound frequency is encoded via circumscribed tuning for sounds of different frequencies, creating a place code for sound frequency (Bulkin & Groh, 2011; De Martino et al., 2013). IC neurons generally are sensitive to both sound location and sound frequency, so both types of codes occur in the same neurons in this structure. Thus, either multiplexing or place coding or both could in principle permit the representation of multiple sounds in the IC.

Testing the “multiplexing” hypothesis required the development of novel statistical methods to detect fluctuating activity patterns and relate them to the expected responses when only one stimulus is present (Caruso et al., 2018; Chen et al., 2026; Glynn et al., 2021; Groh et al., 2024; Marco et al., 2025; Mohl et al., 2020). Using these tools, we have previously found that in the IC, some neurons show patterns consistent with multiplexing when confronted with two sounds that differed in *both* frequency and location (Caruso 2018 ref). As for place coding in the frequency domain, we have found that frequency tuning in the primate IC is very broad (Bulkin & Groh, 2011) and that frequency selectivity degrades in the presence of multiple sounds, suggesting that the place code lacks the resolution to preserve the representation of two perceptually differentiated sounds (Willett & Groh, 2022).

These two previous findings point to multiplexing as a key method of ensuring multiple stimuli can be represented in neural codes, but the relationship between multiplexing and place coding remains uncertain. Does the brain rely on both methods? If so, this would predict that multiplexing should be more prevalent when place coding is least useful, i.e. when the two stimuli in question would not be expected to evoke activity in fully distinct subpopulations of neurons. To test this hypothesis, we evaluated the prevalence of multiplexing in the IC in response to sounds at two different locations across a range of sound frequencies. We found that multiplexing was more common when the two sounds evoked relatively similar response rates, consistent with a greater degree of overlap in the responsive population. This finding suggests that multiplexing is chiefly deployed when place coding “fails” and the stimuli crowd too closely in the neural population map.

## Materials and Methods

### Summary

Data and procedures for the present study were previously published by Willett & Groh (2022). The Duke University Institutional Animal Care and Use Committee approved of all procedures. Single-unit neural responses were recorded from 105 neurons in the IC of two adult female rhesus macaques (Monkey Y right IC, N = 45; Monkey N right IC, N = 59; Monkey N left IC, N = 1) while they localized one or two sounds via saccade. Results are pooled across both monkeys. Each single sound trial consisted of one bandpass filtered noise sound with 10 possible center frequencies (420, 500, 595, 707, 841, 1000, 1189, 1414, 1682, and 2000 Hz) presented either at -12 or 12 degrees from center. Other trials consisted of two bandpass filtered noise sounds with different center frequencies (A and B) presented at the same location (control trials) or 24 degrees apart (dual stimulus trials). There were eight possible A frequencies in 0.25 octave intervals (500, 595, 707, 841, 1000, 1189, 1414, and 1682 Hz) and two possible B frequencies (420 and 2000 Hz). Here, we analyze dual stimulus trials in relation to single stimulus responses. For all the analyses, only correct (rewarded) trials were included. As the monkeys accurately reported the location of each sound during these trials, it is inferred that information about both sounds would be preserved in the neural response, as most auditory inputs must ascend through the IC prior to reaching the auditory thalamus (Aitkin & Phillips, 1984).

### General

We collected behavioral and neural data from two adult female rhesus macaques (Monkey Y – 7 years old at start of recording; Monkey N – 12 years old at start of recording). Procedures have been described in greater detail by Caruso et al. (2018). Eye tracking was accomplished via the scleral search coil method (Judge et al., 1980). The coil as well as a post for restraining head movements were implanted surgically under aseptic conditions and isoflurane anesthesia. After recovery monkeys were trained to complete the sound localization task described in the task section. Once behavioral training was complete, a recording chamber (Crist Instruments) was implanted which allowed access to both the left and right inferior colliculus (IC). However, all but one neuron was recorded from the left IC.

### Stimuli

Auditory stimuli included bandpass filtered noise with center frequencies varying from 420-2000 Hz. The bandwidth of each stimulus was ±10% of its center frequency. Each dual-sound trial consisted of an A frequency and a B frequency. There were eight possible A frequencies in 0.25 octave intervals (500, 595, 707, 841, 1000, 1189, 1414, and 1682 Hz) and two possible B frequencies (420 and 2000 Hz). Each sound stimulus was played with a 10 ms on ramp and an 11 kHz sampling rate. Speakers were located at -12 and +12 degrees horizontally and 0 degrees vertically with respect to the monkey’s head. Speakers were approximately 1 meter away from the monkey. On single stimulus trials, each stimulus was calibrated to 50 dB SPL at the location of the monkey’s head. When combined on dual stimulus trials, the total sound level was 53-55 dB SPL.

### Task

Monkeys localized sounds in three different trial types: single stimulus, dual stimulus, and control. On all trial types, a fixation light appeared to indicate the start of the trial. On attempted trials, the monkey fixated on this light for a variable duration of 600-700ms. Then, in the single stimulus trials, a sound (A or B frequency) was presented either at -12 or 12 degrees from center. Monkeys were required to maintain steady fixation on the central light while this sound played for a variable duration of 1,000-1,100ms. Then, the light would turn off, and the monkey would saccade to the location of the sound. Monkeys would fixate on the location of the sound for 300-650ms to receive a reward.

On the dual stimulus trials, after the initial fixation, rather than a single sound being played at either -12 or 12 degrees from center, two sounds (A and B frequencies) would be played, one at each location. When the central fixation light turned off, monkeys were required to saccade to one of the two locations, hold fixation on this point for 100-500ms, and then saccade to the second location and hold fixation for 100- 300ms.

Control trials followed a very similar structure to single stimulus trials. However, these trials were spectrally equivalent to dual stimulus trials. A combination of two bandpass noise stimulus was played at one location. This ensured that on dual stimulus trials, the monkey was localizing both stimuli rather than simply making two saccades in response to hearing a more spectrally complex sound. Control trials were only completed by Monkey N.

In all trial types, monkeys had a 20 degree horizontal reward window, comparable to other studies using auditory-guided saccades (Metzger et al., 2004). Monkeys were rewarded with a 30% grape juice, 70% water solution. Monkey Y (weight ∼3.7 kg) received ∼0.1mL per trial and ∼60mL per session, while Monkey N (weight ∼9.1 kg) received ∼0.4mL per trial and ∼400mL per session. This provided about 24 attempted trials per stimulus condition per neuron. Further details about the behavioral task are described by Willett & Groh (2022).

### Neural Recordings

A grid and turret system (Crist Instrument, 1mm spaced grid) was inserted into the recording chamber. Grid holes with access to inferior colliculus were determined using structural magnetic resonance imaging (MRI). A single tungsten micro-electrode was inserted into the location of interest, using a stainless-steel guide tube to pierce the dura prior to lowering with a Microdrive (NAN Instruments).

Neural recordings were collected using a Multichannel Acquisition Processor (MAP system, Plexon Inc). Auditory-driven neural activity was detected through listening to single and multiunit signals sent to an external speaker. Advancement was stopped upon identification of auditory-driven activity, and the electrode was allowed to settle for about an hour. Then, the electrode was lowered at 1 *μ*m increments until a single unit was isolated. Spike times from isolated single units (using box method, Plexon SortClient software) were saved for further analysis. Further details about the neural recording are described by Willett & Groh (2022).

### Statistical Analysis

#### Evaluation of Multiplexing

We have previously developed three methods for assessing the presence of multiplexing in neural activity (Caruso et al., 2018; Chen et al., 2026; Marco et al., 2025). These methods differ in how the Bayesian model comparisons are constructed, their reliance on whether neural activity on single stimulus trials meets criteria for being Poisson distributed, the time course of the analysis (across or within trials/spike counting windows), and the role of single stimulus trial benchmarks in model classification. Therefore, each method differs in the sensitivity vs specificity to the presence of multiplexing. However, each can be used to compare the incidence of multiplexing in response to dual-stimulus trials as a function of the difference in individual stimulus responses.

All three models deploy a basic “triplet” structure – the activity evoked by single stimuli (“A” or “B”) is ascertained, and then Bayesian model comparison is conducted to ascertain whether the activity on dual stimulus (“AB”) trials might be caused by some form of fluctuations between (or not) the single-stimulus benchmarks. For all models, only triplets with at least 5 rewarded trials for all three conditions were included, and the neural data was analyzed from 0 to 500ms post stimulus onset.

The original model, here called Spike Count Across Trial (SCAT) (Caruso et al., 2018), determines the fit of the aggregated trial-wise spike count distribution with a mixture of Poisson distributions (switching) as compared to several single-Poisson alternatives. A key precondition for this analysis is that the responses to the two individual stimuli must be discriminable from each other. If the response distributions are too similar, then it would not be possible to ascertain if there is switching between them. Only triplets for which the confidence that the mean spike counts differ exceeds 95% were included for analysis. Note that this criterion limits the range of response overlap that can be investigated in the current study – the responses cannot be identical/fully overlapping.

A second criterion is that the single-stimulus responses must be reasonably Poisson-distributed. Any triplet for which the variance/mean ratio of the single stimulus responses (Fano factor) was greater than or equal to 3 would meaningfully violate this assumption and was excluded from further analysis.

With these criteria in place, the SCAT model compares four possible outcomes, three possible responses that are either ambiguous or reflect alternatives to multiplexing and one response that fits the multiplexing hypothesis. The alternatives are (a) that the responses on dual stimulus trials consistently match one of the two single stimulus distributions, in winner-take-all fashion (“single”), (b) that the responses lie consistently outside the range defined by the two stimulus distributions, e.g. additivity (“outside”), or (c) that the responses lie consistently between the two single stimulus response distributions or represent a weighted average (“intermediate”) – an ambiguous case that could reflect rapid fluctuations occurring frequently within the spike counting window.

Each of these 3 alternatives assumes that the responses will be well fit by a single Poisson distribution; it is only the mean of the distribution that differs across the cases. The fourth possibility is that the AB spike count distribution does not match the shape of a single Poisson distribution but resembles a mixture of the two single-stimulus Poisson distributions. This possibility, “mixture”, is taken as the evidence that the activity on “AB” trials alternates between “A-like” and “B-like”.

The Poisson pre-screening and the benchmarking of the single stimulus response distributions serve as the main safeguards against erroneous classification of mere overdispersion as multiplexing. Nevertheless, this model provides the most generous criteria of the three for identifying fluctuating activity as multiplexing. To limit the potential for false discovery, only cases in which the winning model had a probability greater than or equal to 66.7% - i.e. twice as likely as all other models combined - were included.

Our next model, Spike Count Analysis for Multiplexing Inference (SCAMPI) (Chen et al., 2026), addresses the limitations of SCAT by restructuring the model alternatives. This restructuring allows the detection of multiple forms of fluctuating activity, while retaining competition against single-Poisson options. One category in SCAMPI is “slow switching,” in which spike counts on dual stimulus trials are well fit by a mixture of the two single stimulus Poisson benchmarks (akin to SCAT’s “mixture”). This would detect multiplexing that occurs on a timescale longer than our analysis window (500ms). Another category is “fast switching”, in which the spike count distribution on dual stimulus trials is overdispersed and hence non-Poisson, but remains (stochastically) bounded between the spike count distributions from single stimulus trials. This category is considered sensitive to the possibility that switching may occur within trials (faster than 500ms) such that the spike counts observed on those trials will be distributed across a range roughly bounded by those benchmarks. Additionally, SCAMPI adds an overreaching model, which captures fluctuations in spike count that do not fit as clearly within the single-stimulus benchmarks. These three categories compete against the possibility that the responses are best fit by a single Poisson, whose mean rate may take any value. “Slow switching”, “fast switching” and “overreaching” could all have been classified under SCAT’s “mixture model,” and all three indicate that there is fluctuating activity. With these models competing against each other as well as with the non-fluctuating Poisson alternative, confidence levels were typically lower. Accordingly, triplets for which the winning model had a probability greater than or equal to 50% - at least as likely as all other options combined - were included. As with SCAT, only triplets for which the single stimulus responses are reasonably Poisson and reasonably distinct from each other can be analyzed, so the same exclusion criteria were included.)

Finally, the Multiplexing Activity Regulated by Competition (MARCO) method (Marco et al., 2025) delves more directly into signals at the time scale of individual spikes. MARCO uses Bayesian model comparison to determine the fit of spike times with a competition-based mechanism of multiplexing. Specifically, it proposes that multiplexing-induced spikes arise from competition between two mutually inhibitory drift-diffusion processes, with each encoding one of the presented stimuli. This method disambiguates fluctuations as representing competition between two processes (reflective of multiplexing) or an alternative complex response. This model does not require the Poisson exclusion criteria used for the single stimulus trials for SCAT and SCAMPI; rather, all single stimulus spike-trains must be able to be represented by a time-inhomogeneous inverse Gaussian point process (p > .05). However, as above, the A and B stimuli must evoke a distinguishable response (difference in log pointwise predictive density under one combined model and two separate models for A and B must be greater than log(3)).

These inclusion and exclusion criteria left 199 triplets for SCAT, 293 triplets for SCAMPI, and 311 triplets for MARCO. All statistical analyses were conducted in R.

#### Probability of Fluctuation Classification as a Function of Response Discriminability

We investigated whether an IC neuron’s classification of exhibiting fluctuating activity or not in response to two sound stimuli could be predicted from the discriminability of its responses to each individual stimulus. Triplets classified as “mixture” for SCAT; “slow switching”, “fast switching”, and “overreaching” for SCAMPI; and “slow switching” and “fast switching” for MARCO were all considered to reflect fluctuating activity. Discriminability is measured by *d’_e_*, a measure of sensitivity for signal detection theory for two distributions with unequal variance (Macmillan & Creelman, 2005). Here, measured as the absolute difference in average spike count from 0 to 500ms post stimulus onset divided by the arithmetic mean of the standard deviations of spike count in response to each condition (*d’_e_*, = |*x̅_A_* − *x̅_B_* |/((*s_A_* + *s_B_*)/2)). A mixed effects binomial logistic regression analysis was conducted to determine whether the *d’_e_* of the neuron’s single-stimulus responses affected the probability of that cell being classified as fluctuating in response to the dual-stimulus condition and whether this varied by classification method. This analysis was conducted using the glmer() function in R. The significance threshold for this test was set at α = .05. This model predicted a triplet’s classification as fluctuating or not from the interaction between *d’_e_*and classification method and the nested random effects of cell and stimulus.

## Results

Monkeys performed a sound localization task in which they made saccades to the location of either one (A or B) or two sounds (AB). We used stimuli (bandpass noise with different center frequencies) designed to elicit varying degrees of overlap in the responsive neural population in the IC, with the center frequencies of B sounds flanking those of A sounds. Given that monkeys could successfully perform the behavioral task (∼90% on dual-sound trials compared to ∼80% on single-sound trials, see Willett & Groh, 2022), this suggests that information about both sounds was preserved in the neural response. We propose that when responsive neural populations overlap, time-division multiplexing steps in to preserve these signals. We investigate this claim in the sections that follow.

### IC neurons fluctuate their firing rates in response to two simultaneously presented sounds

First, we analyzed the neural response to dual-stimulus trials (AB) to determine whether neurons switched between “A-like” and “B-like” responses across time. As described in the Methods/Figure 2, we confirmed the presence of multiplexing in this dataset using both the original analysis method (here called Spike Count Across Trial (SCAT) (Caruso et al., 2018), as well as extending the findings using two more recently developed analysis methods: Spike Count Analysis for Multiplexing Inference (SCAMPI) (Chen et al., 2026), and Multiplexing Activity Regulated by Competition (MARCO) (Marco et al., 2025).

**Figure 1.**
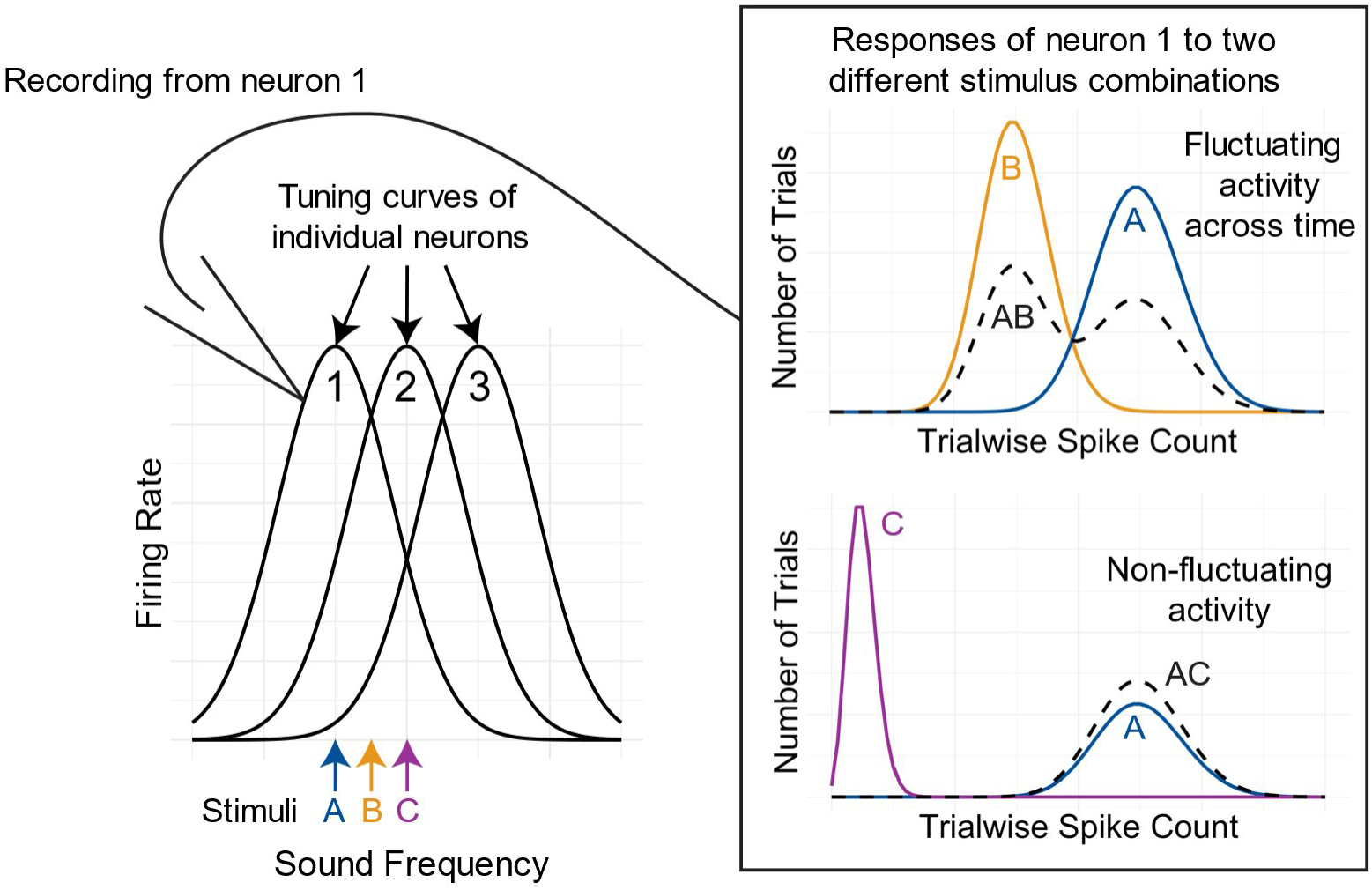
Conceptual link between single-unit responses and resolution of the underlying place code. We expect that as neural responses to each individual stimulus (A and B or A and C in this figure) become more similar, there will be a greater incidence of multiplexing (top right) in response to their combination. Although we are only recording from one neuron at a time (here indicated by the electrode/arrow schematic), the difference in neural response can be thought to reflect the resolution of population code. When two individual stimuli are clearly differentiated by the neural response to each individual stimulus, multiplexing would not be needed. However, when two individual stimuli have more similar responses, multiplexing may be needed in the dual-stimulus condition to preserve information about each individual stimulus.

**Figure 2.**
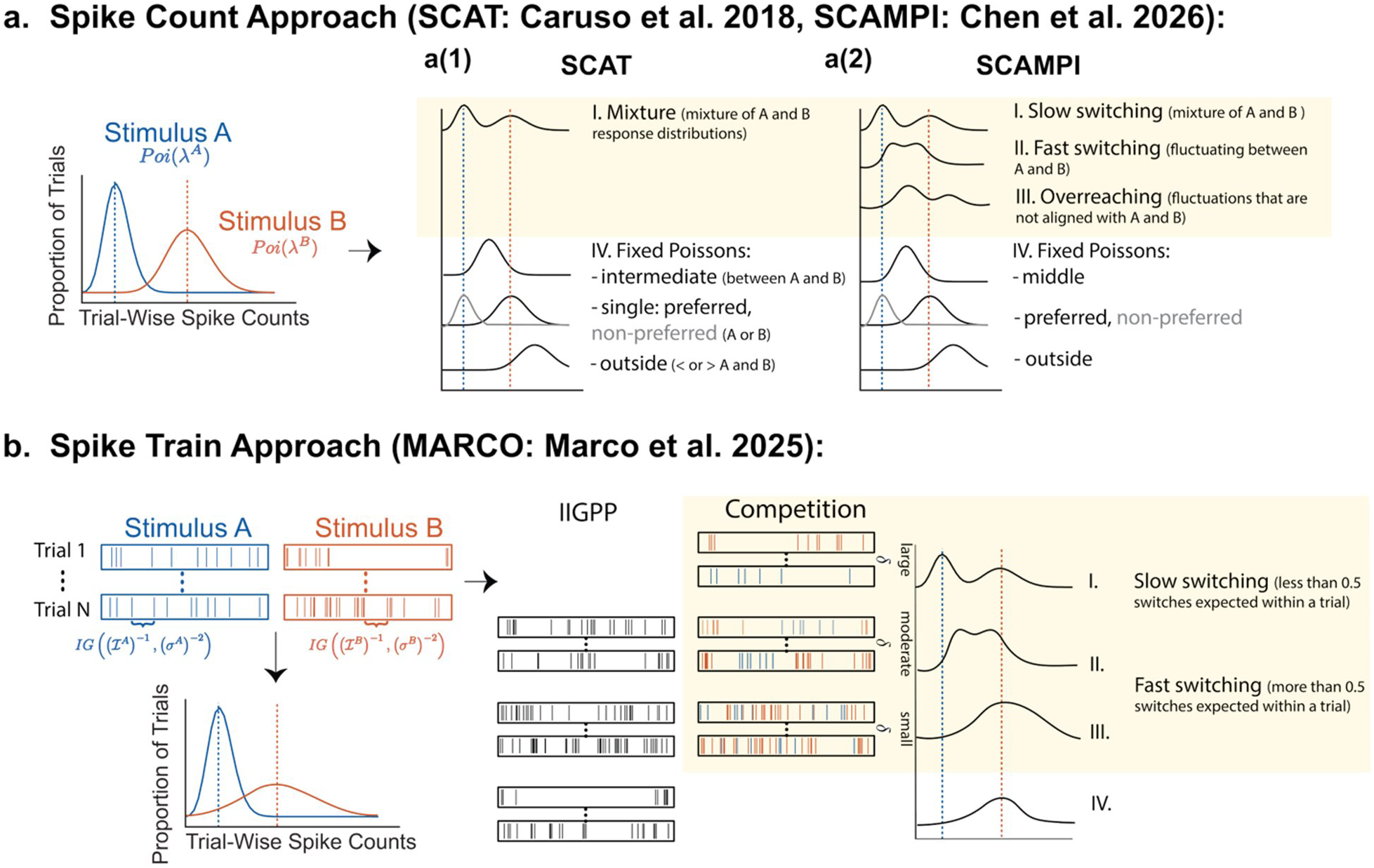
Statistical approaches for detecting fluctuating activity across time. Each approach first characterizes a neuron’s response to two individual stimuli “A” and “B” (left) and then determines whether activity on dual-stimulus trials fluctuates between encoding “A” vs. “B” (right). **a(1).** The original method used to identify multiplexing (here called Spike Count Across Trial – SCAT) compares determines the fit of the distribution of aggregated spike counts in response to the dual-stimulus condition with four possible models, one of which is a combination of the responses to individual stimuli A and B (mixture), and the other three of which represent either consistent, non-fluctuating responses (“single”, “outside”) or an “intermediate” response pattern that could either be non-fluctuating or could indicate response fluctuations that are too fast to be captured by this method. **a(2).** Spike Count Analysis for Multiplexing Inference (SCAMPI) expands upon this method, refining the classification of fluctuating responses into three different competing models: one of which identifies fluctuations across trials/analysis windows (“slow switching”), another of which identifies fluctuations within trials/analysis windows (“fast switching”), and a third possibility that identifies fluctuations that don’t align with single-stimulus benchmarks (“overreaching”). These models compete against a fourth umbrella category of fixed Poissons that encompass the same set of fixed Poisson possibilities as for the SCAT model. **b.** Multiplexing Activity Regulated by Competition (MARCO) takes a different approach. Rather than analyzing aggregated spike counts, MARCO determines the fit of spike times with two competing drift diffusion processes, each encoding “A” or “B”. This model distinguishes between two different types of multiplexing: fewer than 0.5 switches are expected within a trial/analysis window (“slow switching”) or more than 0.5 switches are expected within a trial/analysis window (“fast switching”). This figure is adapted from Chen et al., 2026 and Marco et al., 2025.

Figure 3 presents example neurons identified in the IC to develop an intuition of how the classifications by each of the formal models relate to the observed neural responses. Figure 3a shows the spike count distributions of a neuron in response to a combined “AB” stimulus (gray bars, spline fit) in comparison to the mean spike counts for the “A-alone” and “B-alone” stimuli (orange and blue vertical dashed lines). On the AB trials, there is a bimodal distribution of spike counts, with the modes approximately matching the mean “A” and “B” responses, indicating switching between “A” and “B” like activity patterns. The SCAT model classifies this response pattern as a “mixture”. SCAMPI agrees that this response pattern is consistent with multiplexing but takes note of the fact that the two modes don’t appear to reflect the combination of exactly two underlying Poisson distributions: rather, the spike counts are somewhat spread out between the “A” and “B” means. This could mean that the neuron sometimes switches between the “A” and “B” response categories <u>during</u> the spike counting window, and the model accordingly classifies this response pattern as “fast switching”. Finally, the MARCO method operates on the underlying spike trains (not shown) and uses a definition of “slow switching” if it estimates that fewer than 0.5 switches are expected within a trial under this model. It classifies this particular response pattern as “slow switching” under that definition.

**Figure 3.**
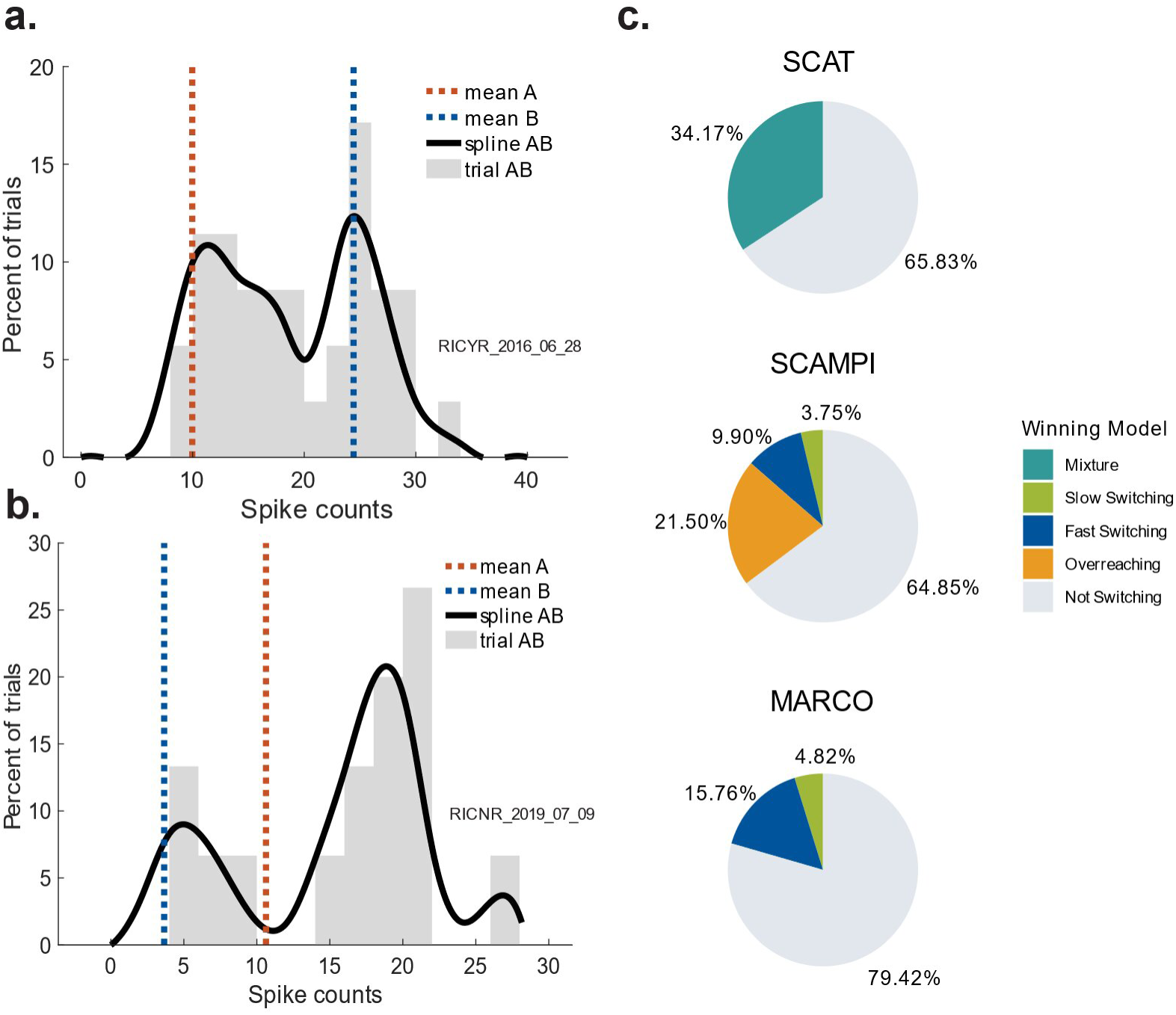
IC neurons can multiplex in response to two simultaneously presented sounds. **a, b.** Two example neurons that both showed bimodal distributions of spike counts across trials. **a.** This neuron (RICYR_2016_06_28) shows fluctuations that are well matched to the single stimulus benchmarks (A: 1189Hz@12 and B: 420Hz@-12) and was classified as a “mixture” by the SCAT model. Note however the presence of spike counts between single stimulus benchmarks – SCAMPI treats these as evidence of switching within trials and classified this as “fast switching”. MARCO evaluated the spike trains and considered the neuron as switching, but with fewer than 0.5 switches per trial, treated it as “slow switching”. **b.** This neuron (RICNR_2019_07_09) showed very evident bimodal fluctuations, but they were not well matched to the single stimulus benchmarks (A: 707Hz@-12 and B: 2000Hz@12); this neuron was classified as “overreaching” by SCAMPI (as well as “mixture” by SCAT and “fast switching” by MARCO). **c.** Percentage of triplets classified as each model of multiplexing compared to the non- multiplexing alternatives. Each model presents evidence for multiplexing in IC neurons.

In contrast, Figure 3b represents a neuron that is less clearly benchmarked between the two component stimulus responses. This neuron is still classified as a mixture under the SCAT method, as it reflects fluctuations in firing rate despite not being as closely constrained to “A-like” and “B-like” responses. The SCAMPI method identifies this distinction, classifying the neuron as “overreaching,” which describes a fluctuating response that does not clearly fit with single-stimulus benchmarks. One possibility for this overreaching classification could be that the two sounds in the dual-stimulus conditions in this dataset are inherently louder than the single-stimulus conditions. This will be addressed further in the discussion. The MARCO method suggests that this neuron’s response reflects competition between two mutually inhibitory drift-diffusion processes, one representing the A stimulus and the other representing the B stimulus. This mutual inhibition is less strong than the previous neuron example, resulting in an expected switch per trial count above 0.5 (“fast switching”). This decreased inhibition may also be the cause of the increased firing rates relative to single-stimulus benchmarks, as it is more reflective of pure competition.

As shown in Figure 3c, a notable proportion of triplets are classified as eliciting fluctuating activity (SCAT: “mixture”; SCAMPI: “slow switching”, “fast switching”, or “overreaching”; MARCO: “slow switching”, “fast switching”), despite not representing the majority of cases. Although triplets may be classified differently by each method given different assumptions (see Figure 3a), we see a similar proportion classified as fluctuating across methods, with slightly fewer under the MARCO model given that it has the strictest definition of multiplexing. Next, we investigated what could be driving neurons to multiplex and whether it depended on the resolution of the neural code in response to single stimuli.

It is beyond the scope of the present paper to make claims about which model is “best”. Rather, we are interested in how the proportion of multiplexing evident in any given method relates to the difference in responsiveness evoked by the “A” stimulus vs the “B” stimulus.

### The populations of neurons responsive to each stimulus are largely overlapping

Similar to previous reports with tones (Bulkin & Groh, 2011), we find that neural tuning to bandpass noise is sufficiently broad such that a large proportion of the recorded population is capable of responding to any particular stimulus. Figure 4 quantifies the proportion of neurons responsive to each of our individual stimuli across the neural population we recorded. We considered a neuron to be responsive to a given single stimulus if its response 0-500 ms after stimulus onset was greater than baseline (-500:0 ms prior to stimulus onset) using a one-sided paired t-test (α = .05, uncorrected). This identifies cells that are excited by the stimulus. The proportion of the total population responsive to each A stimulus regardless of presented location is shown in Figure 4a and ranges from 46-86% of the population.

**Figure 4.**
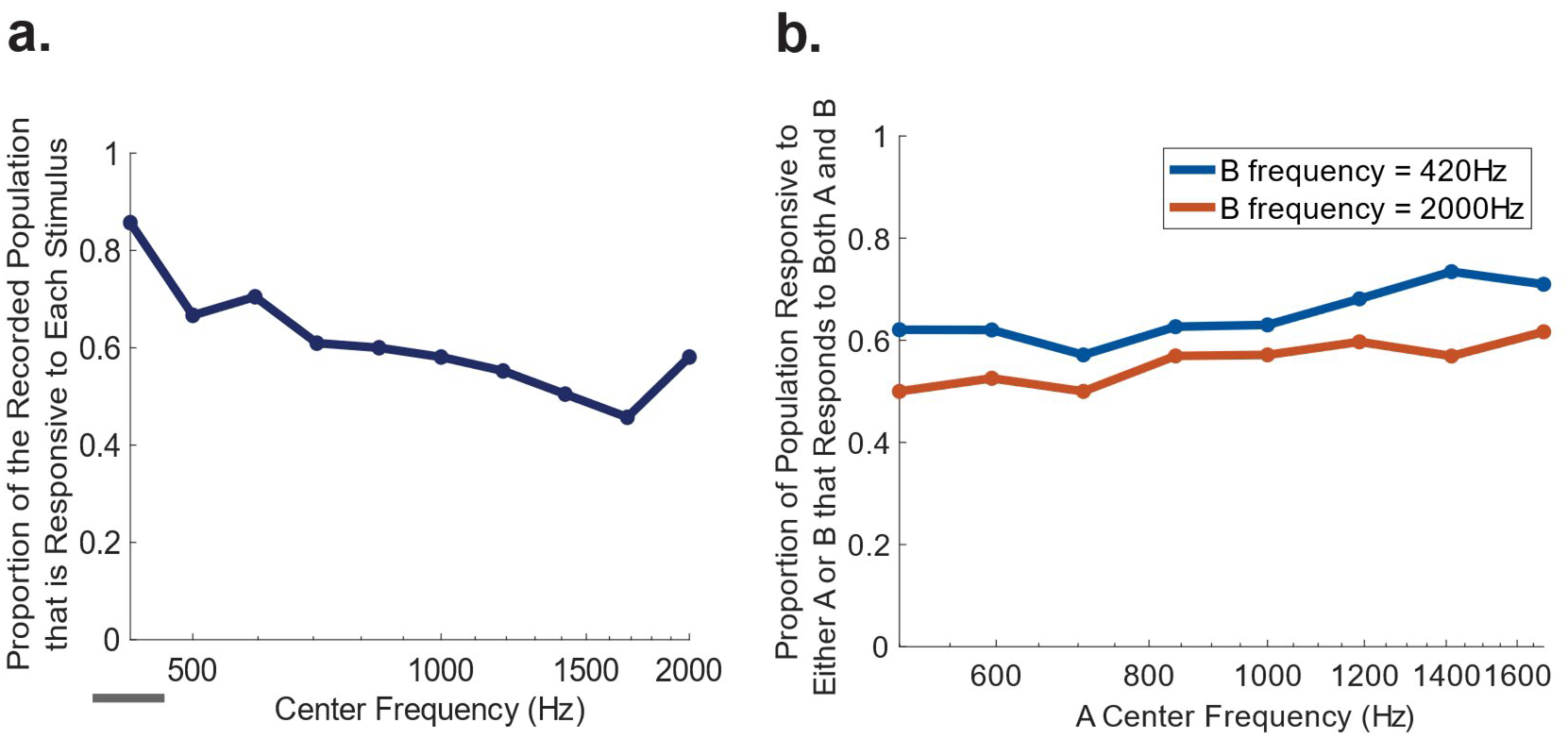
The population of neurons responsive to each stimulus is largely overlapping. **a.** The proportion of neurons recorded from that are responsive to each stimulus depending on the center frequency regardless of location presented. The gray bar below the figure represents the bandwidth. The neural population is more responsive to lower frequencies, but at least 46% of the population responds to each stimulus. **b.** The proportion of neurons responsive to either stimulus that are responsive to both stimuli is displayed for each frequency pairing presented (different location pairs only). Even for relatively large frequency differences, the percentage of population overlap does not drop below 50%.

Next, we calculated the number of cells that were responsive to both stimuli within each stimulus pair and divided this by the total number of cells responsive to either stimulus (Figure 4b). We did this for only A and B sounds at different locations, as it was shown behaviorally that monkeys could distinguish between these pairs (see Willett & Groh, 2022). Depending on the “B” stimulus’ frequency, 50-62% of neurons exhibited significant responses to both individual stimuli that could be combined in the AB conditions.

In short, within our sample, we find that there is a considerable amount of overlap in the neural populations responsive to each stimulus, even for stimuli with greater frequency differences (see Figure 4b). Given this, and given that multiplexing was present in some neurons/stimulus conditions and not others, we can ask whether the similarity in responsiveness to a given pair of stimuli predicts the likelihood of multiplexing being observed.

### As responses to component stimuli become more similar, there is a greater incidence of multiplexing

To determine whether the incidence of multiplexing depends on the discriminability of individual stimulus responses, we calculated the *d’_e_*of the neural response to the two single-sound conditions (Macmillan & Creelman, 2005) for each triplet. This was the absolute difference in average spike count from 0 to 500ms post stimulus onset divided by the mean of the standard deviations of spike count in response to each condition (*d’_e_ =* |*x̅_A_* − *x̅_B_* |/((*s_A_* + *s_B_*)/2)). This measure captures both the effects of stimulus similarity and the neural response properties (e.g., tuning bandwidth) on the ability of a neuron to distinguish between the two component stimuli.

To visualize the relationship between response *d’_e_* and incidence of multiplexing, we binned each triplet into one *d’_e_* bins and calculated the proportion of triplets in each bin classified as exhibiting a fluctuating response across each of the three models. For the SCAT model, this included triplets classified as mixture. For the SCAMPI model, this included triplets classified as fast switching, slow switching, and overreaching. For the MARCO model, this included triplets identified as fast or slow switching.

Qualitatively, as the neural response to each single-stimulus condition in the triplet became more similar, there was a greater incidence of fluctuating activity (see Figure 5). To quantify this formally, a mixed effects binomial logistic regression analysis was conducted to determine whether the *d’_e_* of a neuron’s response to the two single-stimulus conditions affected the probability of that cell being classified as multiplexing in response to the dual-stimulus condition and whether this effect depended on the method used to classify the response. As shown in Figure 6, we found a significant main effect of *d’_e_* (z = -2.73, p = .006), with the probability of multiplexing decreasing with greater *d’_e_*. We found no significant interaction effect, suggesting that this effect did not significantly differ among the models used to classify multiplexing responses. However, we did find a significant main effect of the MARCO model on the probability of multiplexing (z = -3.06, p = .002). This suggests that the MARCO model classified significantly less triplets as multiplexing compared to the reference SCAT model. This is to be expected, as MARCO has a stricter mechanistic definition of multiplexing.

**Figure 5.**
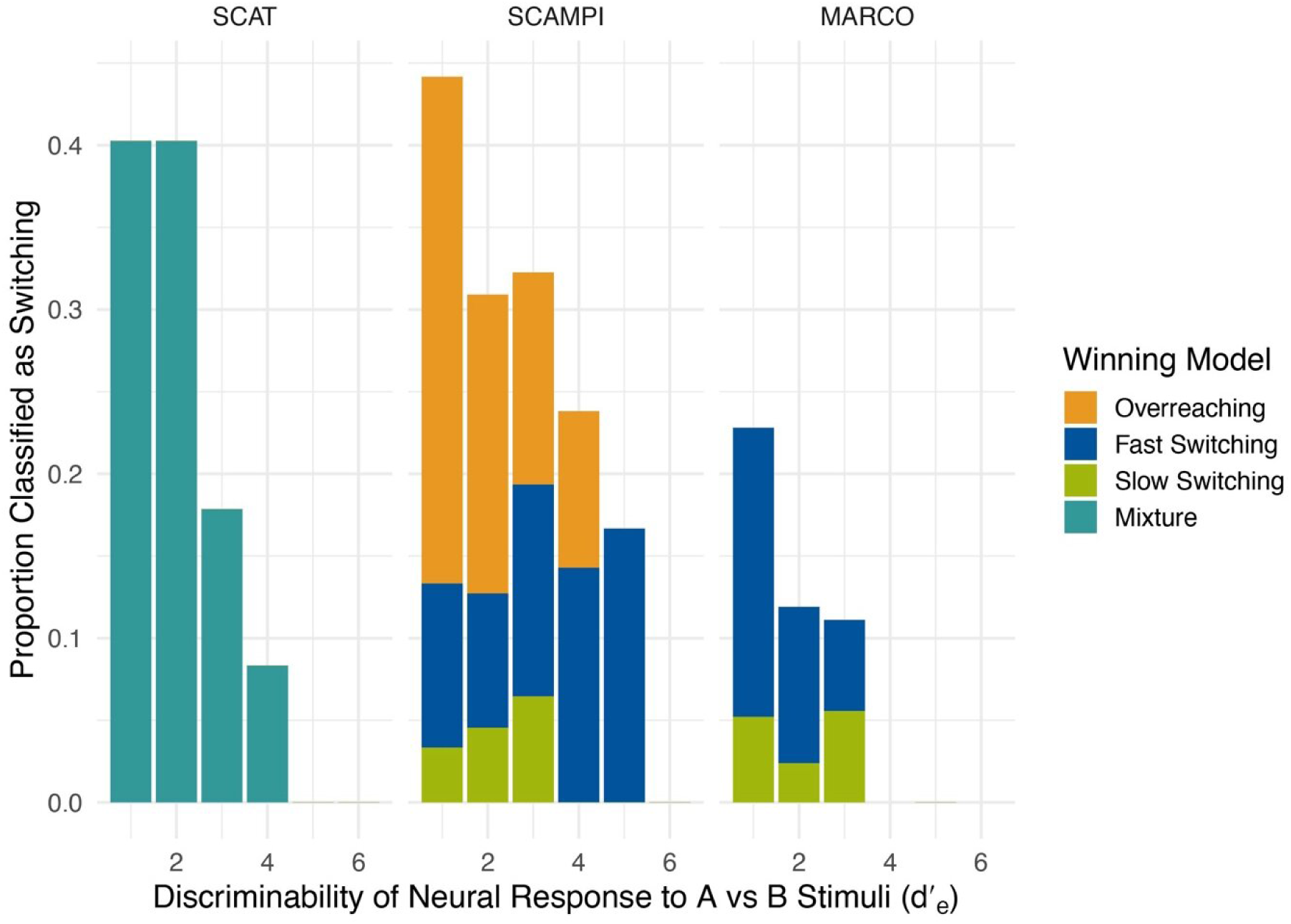
As responses to component stimuli become more similar, there is a greater incidence of multiplexing. The proportion of triplets classified as each model of multiplexing is displayed across each d’_e_ bin. The trend is the same for each of the three models used to detect time-division multiplexing.

**Figure 6.**
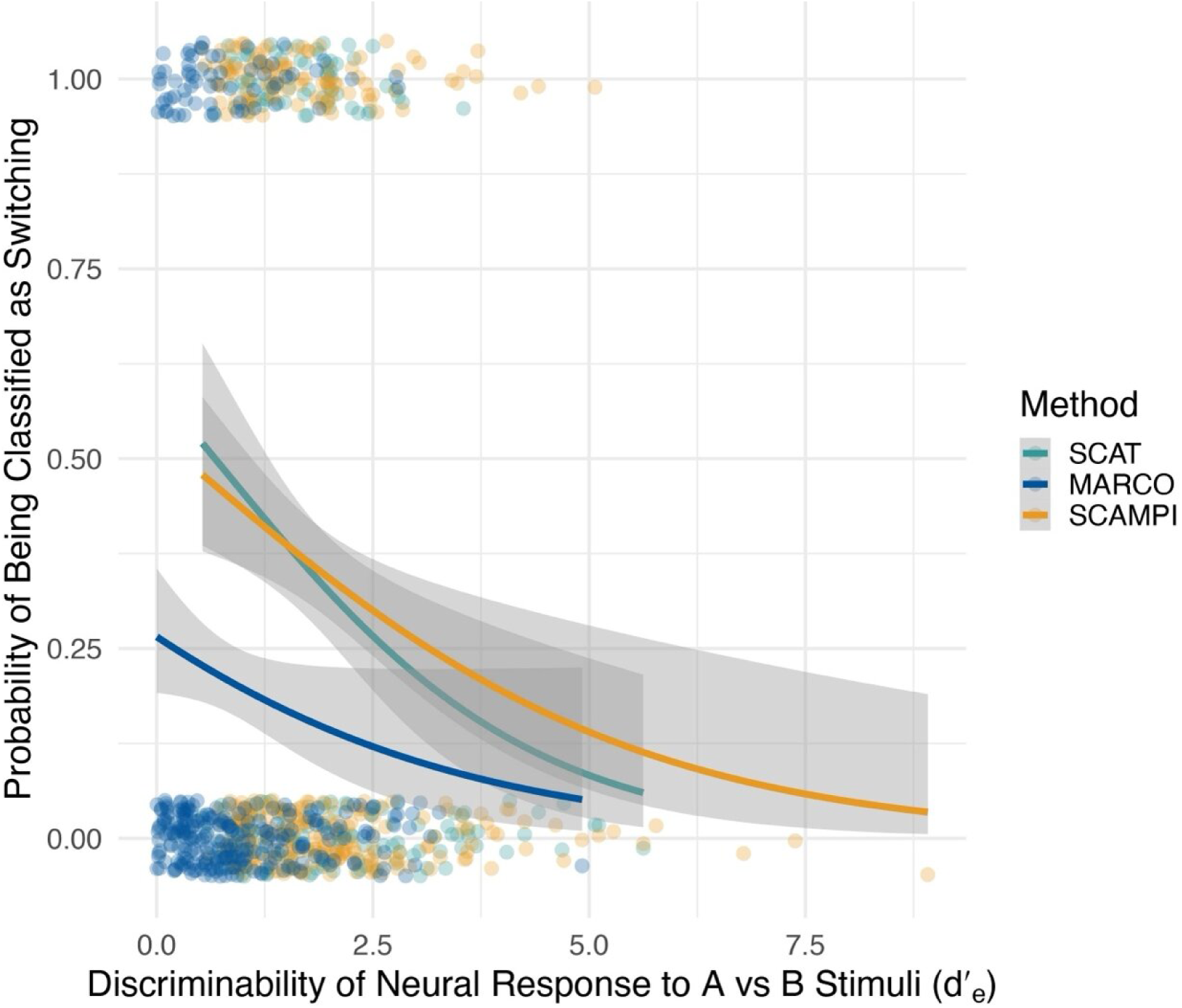
Neurons have a greater probability of multiplexing in response to two stimuli for which the response is more similar. A mixed effects binomial logistic regression analysis was conducted to determine whether the discriminability of the neural response to two individual stimuli A and B affected the probability of a cell multiplexing in response to their combination and whether this effect differed across classification methods. This model predicted the binary classification of multiplexing (1) or not multiplexing (0) for each triplet, displayed as individual data points on the plot (with some vertical jitter to visualize the density of samples) from the interaction between d’_e_ and statistical method with random effects of cell and stimulus condition nested within cell (triplet), multiplexing classification ∼ d’_e_ * statistical method + (1 | cell / stimulus). Neurons were significantly less likely to be classified as a mixture response to dual-stimulus conditions when they had a greater d’_e_ to the matching single-stimulus conditions (z = - 2.73, p = .006). The interaction between d’_e_ and statistical method was not significant; however, the MARCO method was significantly less likely to identify a triplet as multiplexing (z = -3.06, p = .002).

### Individual neurons can multiplex under some stimulus conditions and not others

The dependence of multiplexing on stimulus conditions in the previous section implies that at least in some cases, the same neuron may multiplex to one set of stimulus conditions but not to another. Figure 7 demonstrates this across the neural population, using the SCAT approach. Each individual neuron is plotted as a row/connected set of dots on the y axis, and each dot shows the “mixture” probability for a particular set of stimulus conditions for that neuron. Only conditions that passed criteria for analysis are illustrated, so most neurons do not have dots for every stimulus condition that was tested. Neurons are sorted according to their average mixture probability across stimulus conditions.

**Figure 7.**
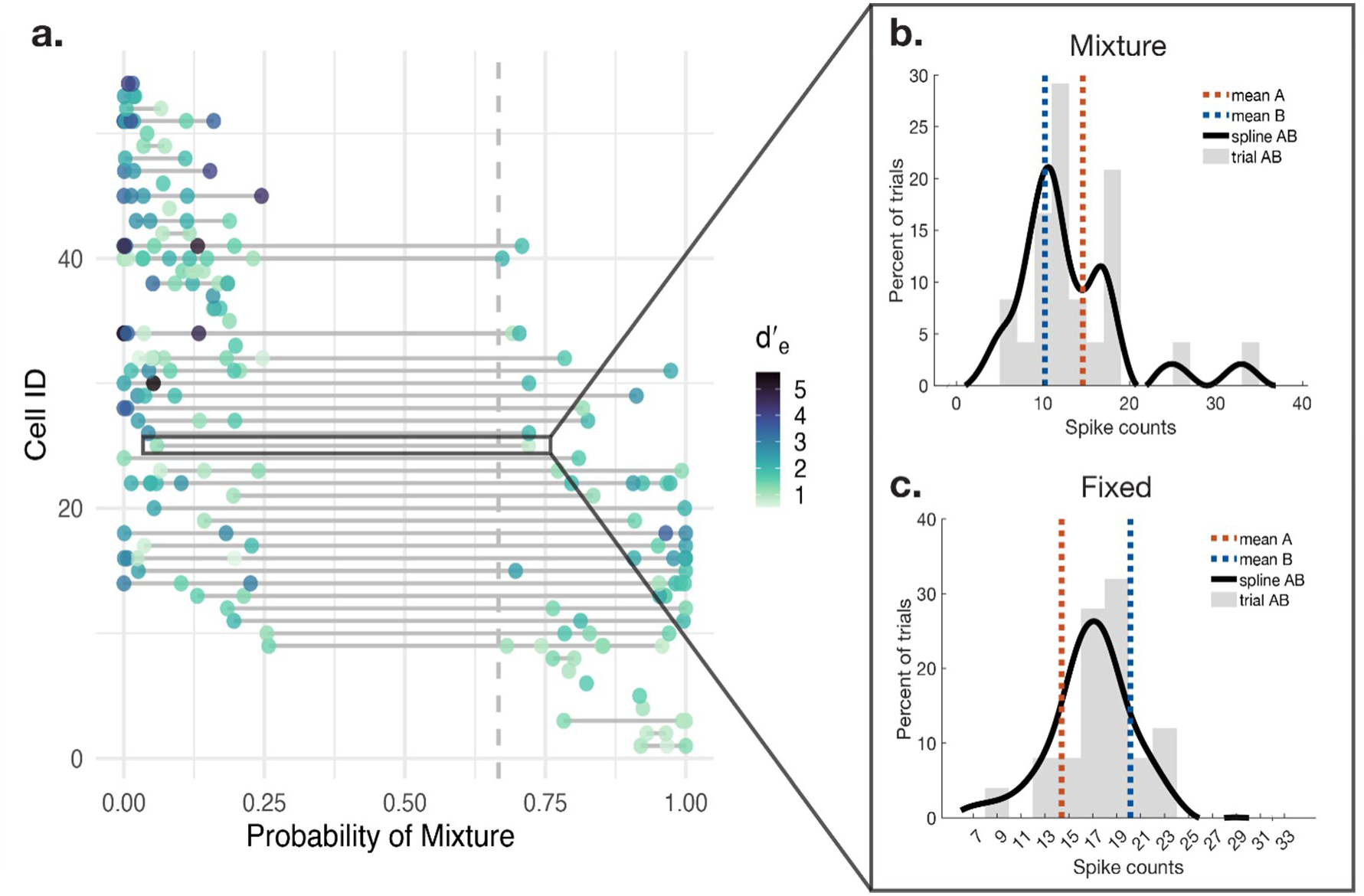
Multiplexing depends on both the cell and the stimulus. **a.** The probability of the dual stimulus response fitting with a mixture model is displayed for each cell across conditions under the SCAT classification method. Each cell is represented as a horizontal line, with that cell’s response to a particular triplet represented by a colored circle. The color of the circle represents the cell’s difference in neural response to component stimuli, with darker circles representing a greater difference in neural response. The vertical dashed line represents the winning probability threshold for multiplexing classifications. Only triplet conditions that pass the exclusion criteria for the SCAT method are included. Many neurons vary in whether they multiplex depending on the stimulus condition. **b, c.** The panel to the right is an example of a cell (RICYR_2015_12_16) that multiplexes in response to one stimulus condition (**b**, A: 707Hz@12, B: 2000Hz @-12) and not another (**c**, A: 1000Hz@12, B: 420Hz@-12).

Across all cells in the neural population recorded from, we find that a majority of cells multiplex in at least one condition (64.8%): about two thirds of the neurons plotted in Figure 7 exhibited at least one point to the right of the dashed line. Consistent with the analyses in the previous section, these conditions tended to have lower or intermediate *d’_e_*values (the corresponding dots have lighter blue colors). At the same time, a majority of neurons also exhibited response patterns that were not well classified as “mixtures” in at least one condition (85.2%, data points to the left side of Figure 7); these included the dots that had high *d’_e_* values (darker blues) indicating large response differences between the A and B stimuli. The middle range of this graph highlights neurons that showed “mixture” responses to some stimuli and not to others.

Altogether, this analysis shows that the ability to multiplex is both stimulus dependent and dependent on the individual neuron: some but not all neurons appear to have the ability to multiplex if the conditions are right.

## Discussion

Here, we demonstrate that as the neural responses to two individual stimuli become more similar, neurons are more likely to multiplex in response to their combination. This suggests that rather than relying on either multiplexing or place coding alone to preserve information about two simultaneous stimuli, the brain relies on both, recruiting multiplexing when it is most needed due to coarseness of the neural place code. These findings provide insight into the role of multiplexing within the broader framework of neural coding, suggesting that place coding and multiplexing may serve as complementary mechanisms.

Specifically, we confirmed that tuning in the IC was sufficiently coarse that the neural populations responsive to two of our auditory stimuli varying in frequency and location were largely overlapping. This population overlap was associated with a propensity to multiplex, with the probability of a neuron multiplexing increasing as that neuron’s response to each individual stimulus became more similar. This effect did not significantly differ across three different multiplexing classification methods, adding to its robustness.

Each of these methods differ in their sensitivity and specificity as well as their mechanistic interpretability. Of particular ambiguity is the overreaching model classification under the SCAMPI method. The majority of switching classifications by the SCAMPI method fell under this category (see Figure 3c). Fit with this model suggests that fluctuations across time are not clearly aligned with single-stimulus benchmarks. However, this model is agnostic to how these fluctuations deviate from the single stimulus responses and the reason for this overdispersion.

A possible explanation is that the loudness of the dual stimulus conditions was greater than the single stimulus conditions (due to combination of the two sounds). This increase in loudness could raise the overall firing rate. Thus, the firing rates that the neuron fluctuated between might have been shifted with respect to that of the response to the component stimuli. This reveals the importance of ensuring dual stimuli reflect a true combination of individual stimuli in future analyses.

An overreaching classification could also reflect the effect of competition with low inhibition between the two processes encoding each signal to elicit a spike, causing an increase in the overall firing rate. Imagine, for example, that you have two entrances to your home that both connect to the same doorbell tone. Guests arrive at each of these entrances independently, but the first bell to be pressed elicits the doorbell tone. Even though each entrance is independent, the doorbell will be more likely to ring earlier since the faster of the two doorbell presses elicits the tone. The same could occur with two independent neural signals competing to elicit a spike, and this would also be reflective of a multiplexing signal.

However, an overreaching classification could also reflect non-information-preserving noise. Rather than switching between each stimulus, there could be non-stimulus related signals that vary from trial to trial, impacting the firing rate of the neuron. This would not fit with a multiplexing response.

This ambiguity is addressed with the MARCO method, demonstrating the benefit of including complementary classification methods. MARCO directly tests whether the neural spike times fit with a competition-based mechanism of multiplexing. This method identified overall fewer triplets as multiplexing, as it has a stricter definition. However, there was no significant difference in the relationship between response discriminability and probability of multiplexing classification between MARCO or SCAMPI and the SCAT method (Caruso et al., 2018; Chen et al., 2026; Marco et al., 2025), suggesting that this relationship holds with a more specific analysis.

Although we do see a relationship between response discriminability and multiplexing classification, even when responses to individual stimuli become very similar, multiplexing remains the minority response to the dual stimulus. Therefore, we do not claim that all neurons multiplex in response to two stimuli with overlapping representations in the neural population; rather, we suggest that some neurons multiplex, and such neurons are more likely to multiplex when their response to each individual stimulus is less distinct. This highlights the importance of viewing multiplexing as one mechanism within the overall population code and motivates the need for future research investigating how multiplexing is coordinated across a neural population, leveraging simultaneous recording methods.

Although we do not currently have information about simultaneously recorded neurons, we can begin to consider how multiplexing might be organized across a neural population based on the current results. When two stimuli activate distinct neural populations at an early processing level, fluctuating activity may not be needed to distinguish the responses to each individual stimulus. However, at a further level of processing, there may be overlap in the neural populations responsive to each stimulus due to convergence. To coordinate multiplexing, these populations could simply rely on short inhibitory connections between neighboring neurons with distinct tuning preferences that converge onto the same downstream populations (see Figure 8).

**Figure 8.**
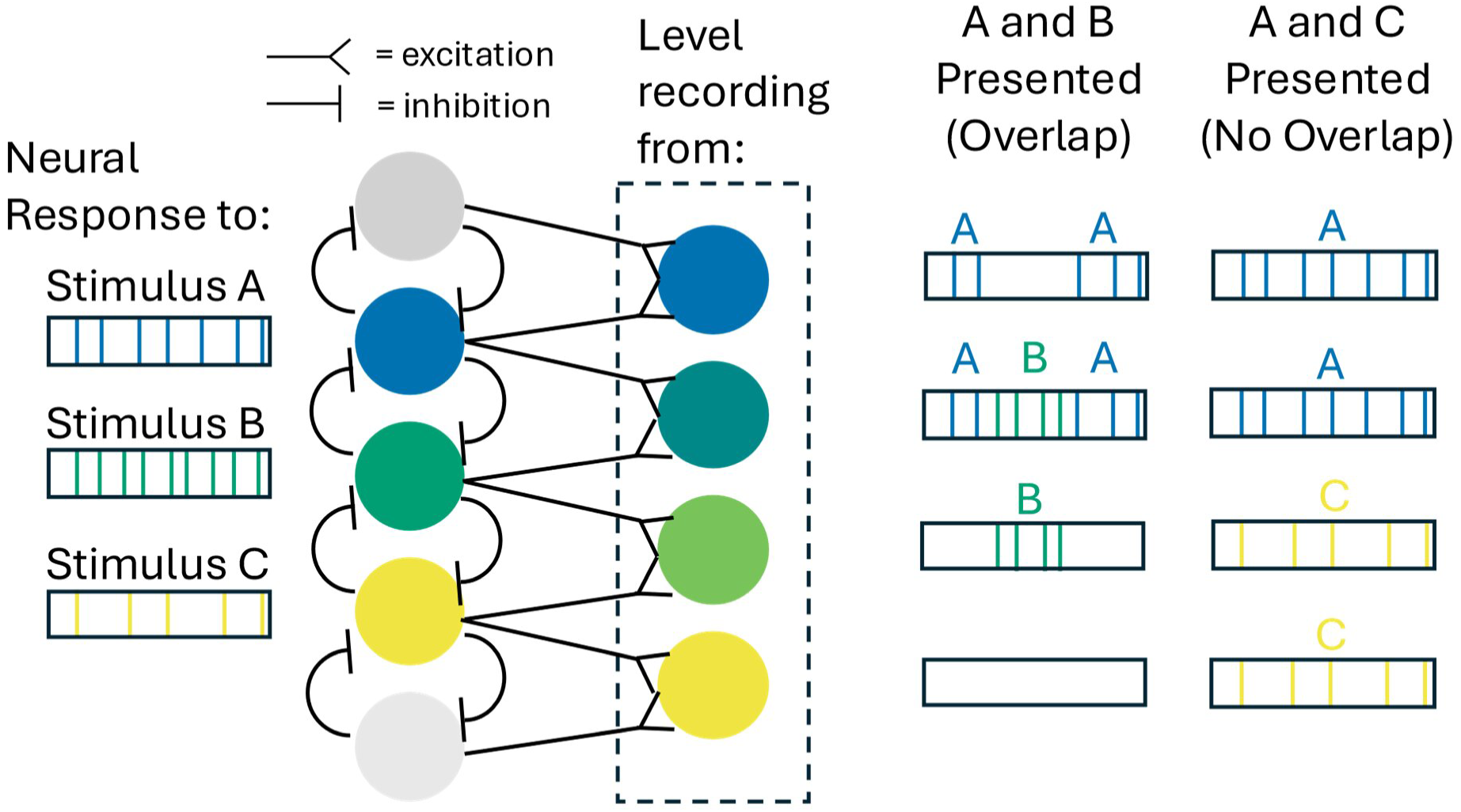
Local connections may be sufficient to coordinate multiplexing. In the first column of neurons (left), there is no overlap in the populations responding to each stimulus. Thus, multiplexing would not be required at this level. However, these neurons converge on a downstream population (column to the right), causing overlap in the neural populations responsive to stimuli A and B and B and C (with no overlap between stimuli A and C). As shown in the figure, local inhibitory connections between neighboring neurons with different stimulus preferences would be sufficient to generate multiplexing in the neurons that respond to both stimuli (A and B), and they would not generate multiplexing for stimuli with no population overlap (A and C). Figure adapted from Marco et al., 2025.

Under this model, top-down input or long-range connections across populations that do not have overlapping outputs would not be needed. This model could realistically be implemented in the inferior colliculus, as sideband inhibition – when a neuron inhibits a neighboring neuron with slightly different tuning preferences – has been suggested as an essential mechanism that shapes tuning curves in the IC (Pollak et al., 2011).

Therefore, multiplexing could emerge from existing connections in the IC as well as other sensory regions (Born & Tootell, 1991; Liu & Kanold, 2021).

## Acknowledgments

This work was supported by NIH grant R01 NS129112 to JMG and STT. We are grateful to Stephanie Lovich and Gelana Tostaeva for laboratory assistance.

